# From rest to focus: Pharmacological modulation of the relationship between resting state dynamics and task-based brain activation

**DOI:** 10.1101/2025.09.30.679567

**Authors:** Kathryn Biernacki, Tianye Zhai, Justine Hill, Emily McConnell, Betty-Jo Salmeron, Roselinde Kaiser, Amy Janes

## Abstract

Dynamic resting-state brain activity provides insight into intrinsic neural function and holds promise for predicting individual responses to cognitive demands and pharmacological interventions. This research could ultimately guide medication selection, yet links between network dynamics and medication effects on cognitive function require further validation. Here, we examined whether dynamic activity of an attentional network at rest relates to task-evoked brain activation on the Multi Source Interference Task (MSIT) following administration of methylphenidate (20mg) and haloperidol (2mg), which have opposing effects on attention and catecholaminergic function. Fifty-nine healthy adults completed resting-state and task-based fMRI on three separate days on which they received methylphenidate, haloperidol, or placebo in a double-blind placebo-controlled design. Coactivation pattern analysis determined time spent in the dorsal attention network (DAN) under placebo at rest. Linear mixed-effects modelling assessing the relationship between MSIT task activation under drug and time spent in DAN at rest under placebo and MSIT task activation under drug identified a significant interaction in the dorsolateral prefrontal cortex (dlPFC; p<0.001). Post-hoc analyses indicated that more time in the DAN at rest under placebo was associated with decreased MSIT dlPFC activation under methylphenidate and increased dlPFC activation under haloperidol. Findings demonstrate that resting dynamics of an attentional network are linked to task-related brain responses under different drug conditions within a region implicated in attentional control and sensitive to catecholaminergic variance. Resting-state dynamics may predict pharmacological modulation of goal-directed cognition, highlighting the potential clinical utility of resting-state dynamics in predicting medication response and supporting individualized treatment.

## Introduction

Resting-state functional magnetic resonance imaging (rs-fMRI) is a powerful tool for studying the brain’s baseline function. It provides valuable insights into the neural correlates of undirected cognition that are also pertinent to task-evoked activity: substantial evidence links resting-state brain function with subsequent task-based activation [1–6]. Indeed, it has been suggested that common mechanisms drive brain activation at rest and during task performance, allowing for the prediction of task performance based on rs-fMRI [7,8]. Critically, rs-fMRI also provides valuable insights into the neurobiological impact of pharmacology [9–12] and treatment outcomes [13–16], including the ability to predict the impact of commonly prescribed psychiatric medications [17–20]. Taken together, previous work suggests that rs-fMRI may be a clinical tool in predicting how medication administration will impact cognition and sets the stage for examining how rs-fMRI, in drug-free conditions, may be used to understand how medication administration will impact brain activity during task-directed cognition. To investigate these ideas, the present work investigates how brain function at rest, under drug-free conditions, informs pharmacological response to attention-altering medications during cognitive demand.

Previous work linking resting state with task-based activity and pharmacological response has traditionally used static approaches to measuring resting brain connectivity. Static methods typically capture brain functional connectivity across the entire fMRI scan [21,22]. While static methods of analyses provide valuable insights into the brain’s functional organization, the human brain operates dynamically, including during “rest” [22]. Dynamic methods provide insights into brain function and organization by measuring time-varying changes in brain connectivity or activity, where individual differences in such temporal properties are tied to individual differences in cognitive ability [21,23,24]. One analysis approach used for evaluating dynamic properties of brain networks, co-activation pattern (CAP) analysis, identifies recurring patterns of brain activation [24]. This method enables the examination of temporal properties of transient network states, including "time-in-state", which is a measure of how long the brain spends in a particular network state. In a large group of healthy individuals, we previously showed that the CAP method defined transient brain states which spatially overlap with well-defined resting state networks such as the dorsal attention network (DAN), default mode network (DMN), salience network (SN), and others [25]. Further, the temporal properties of these networks replicated across independent scanning days, showing that network dynamics are relatively stable. Our group later showed that resting network dynamics were correlated with task-related brain activation [6], suggesting that time spent in specific resting states may relate to how the brain responds during subsequent tasks. Moreover, [10] demonstrated that that improvements in depressive symptoms following sertraline treatment were mediated by dynamic function of CAP-derived DMN and SN states, suggesting that dynamic functioning of brain states may correspond with medication response. Crucially, time in state is biologically relevant [21,26] and has also been shown to predict variability in task-related brain activation [6], is sensitive to pharmacological manipulation [27,28] and differs between clinical groups [29–31]. Collectively, these findings suggest that temporal dynamics, particularly time in state, can be used to assess links between resting brain function, task-related responses, and medication effects. However, the specific role of pharmacology in modulating the link between resting dynamics and task-related activity has not been directly studied. This study addresses this gap by examining how medication-free resting brain dynamics relate to task activation under different pharmacological conditions.

In this study we evaluated how resting brain dynamics under placebo interact with pharmacology to relate to task-based brain activation in the context of attentional processing. We assessed task-based brain activation during the multisource interference task (MSIT), a well-established cognitive control task that robustly activates attention-related brain regions [32–34]. Methylphenidate and haloperidol were used to modulate pharmacological state during the task, as both medications affect catecholaminergic neurotransmitters and cognition in opposing ways. Methylphenidate, a medication for attention deficit hyperactivity disorder (ADHD), increases dopamine and norepinephrine by blocking reuptake of these neurotransmitters, leading to enhanced processing speed, overall and sustained attention [35,36]. In contrast, haloperidol, a dopamine D2/D3 receptor antagonist, slows reaction times and reduces performance on cognitive tasks requiring sustained attention [37,38]. The impact of these pharmacological interventions on attention are likely linked to the modulation of networks that support performance of cognitive tasks that place demands on external attention. Previous work has shown that methylphenidate increases attention task-based brain activity [39–42], and while at rest, methylphenidate impacts network connectivity in a manner that contributes to methylphenidate’s attention-enhancing effects [43–45]. Conversely, haloperidol has been shown to attenuate brain activation in attention networks during attentional tasks [46,47] and at rest [48,49]. In our previous work we also demonstrated that methylphenidate and haloperidol modulated the temporal dynamic properties of large-scale cognitive networks at rest [27]. However, what is unknown is how drug-free temporal dynamics at rest relate to task-based brain activation under drug. As such, we assessed attention-related brain activation as a function of the interaction between drug (methylphenidate and haloperidol) and the resting-state dynamics of the dorsal attention network (DAN) during the placebo condition. We focused on the DAN as this network is involved in externally oriented attention [50], where the activity of this network is selectively increased during tasks that place high demand on externally focused attention [51] and its activity is tightly tied to the function of catecholaminergic neurotransmitters such as dopamine [52]. Moreover, the involvement of the DAN in cognitive processes is highly consistent across rest and task [45]. Together, this makes the DAN an ideal probe for understanding how dopaminergic medications known to impact attention-related cognition are linked to resting dynamics. Measuring how the engagement of the DAN under placebo is related to task-based activity under methylphenidate and haloperidol provides another step in understanding how resting brain function may be used to predict pharmacological response.

## Materials and Methods

### Participants

Partial data from this participant sample have been previously published in Hill et al., 2024 [27]. Participants included 59 healthy right-handed individuals (see Table 1 for participant characteristics). Participants were excluded for: meeting DSM-IV diagnostic criteria for lifetime substance dependence and/or substance abuse in the 2 years preceding the study; current or history of neurological illness or major psychiatric disorders (such as schizophrenia or bipolar); cognitive impairment; cardiovascular conditions; health conditions or use of medications that interfere with the BOLD signal or alter metabolism of catecholaminergic agents; and contraindications for MRI. Participants were recruited from Baltimore, MD, and surrounding areas. The study was a registered clinical trial (NCT01924468). The study was approved by the National Institute on Drug Abuse-Intramural Research Program Institutional Review Board and written informed consent was obtained from each participant.

**Table 1.** Participant characteristics.

| Demographics |  |
| --- | --- |
| Age (years) | 30.97 ($SD=9.42$) |
| Education (years) | 15.41 ($SD=3.05$) |
| Sex (male/female) | 17/42 |
| Race (*n*) |  |
| Black or African American | 11 |
| White | 37 |
| Asian | 5 |
| More Than One Race | 6 |
| Ethnicity (*n*) |  |
| Hispanic or Latino | 9 |
| Not Hispanic or Latino | 49 |
| Unknown or Not Reported | 1 |

### Experimental Procedures

All participants received an acute oral dose of 20 mg methylphenidate/ 2 mg haloperidol/placebo on 3 separate study visit days in a double-blind randomized order. fMRI scans took place at drug peak for each drug condition: 4-hours post haloperidol/placebo and 1-hour post placebo/methylphenidate; timed in accordance with absorption rates to ensure high and stable plasma levels of medication during the scan. See Supplementary Figure 1 for a schematic of the timing of drug administration and scan schedule. All study days were identical.

### Imaging procedures

All participants completed a resting state fMRI scan followed by a task-based fMRI scan (see Supplementary Figure 2). For the 8-minute resting state scan, participants were instructed to keep eyes closed but stay awake. For the 11-minute task-based fMRI, participants completed an event-related multi-source interference task (MSIT). In this task, subjects were instructed to identify, via button-press in the scanner, a target number among the three numbers presented on screen. Specifically, subjects were instructed that the target number presented on screen would always be different than the other two numbers. The numbers displayed on the screen match the buttons, which are arranged in the same left-to-right linear order. If the target number was “1” participants were instructed to press the corresponding first, left most, button on the button box for “2” the second, middle, button and for “3” the third, right most, button. On congruent trials, the target number matched the position on the button box (e.g., “1” in the leftmost position on screen and the number “0” was in the second and third place on the screen); on incongruent trials, the target number never matched its position on the button-box (e.g., “1” in the rightmost position on screen and another number such as 2 or 3 is in the first and second position on the screen – for this trial the subject has to inhibit responding to the number location (the third space) and instead has select the first leftmost button). Participants completed one run of the MSIT comprised of three blocks with 24 trials each. Congruent and incongruent trials were presented in random order within each block with a 50% split between trial types (12 congruent, 12 incongruent). Stimuli were presented for 3s and inter trial intervals (ITIs) were randomly jittered from 4 to 8s in 500ms increments. Full task details of the block-design MSIT are reported in [32]. See Figure 1 for a schematic of the event-based version of the task used in the current protocol.

**Figure 1.**
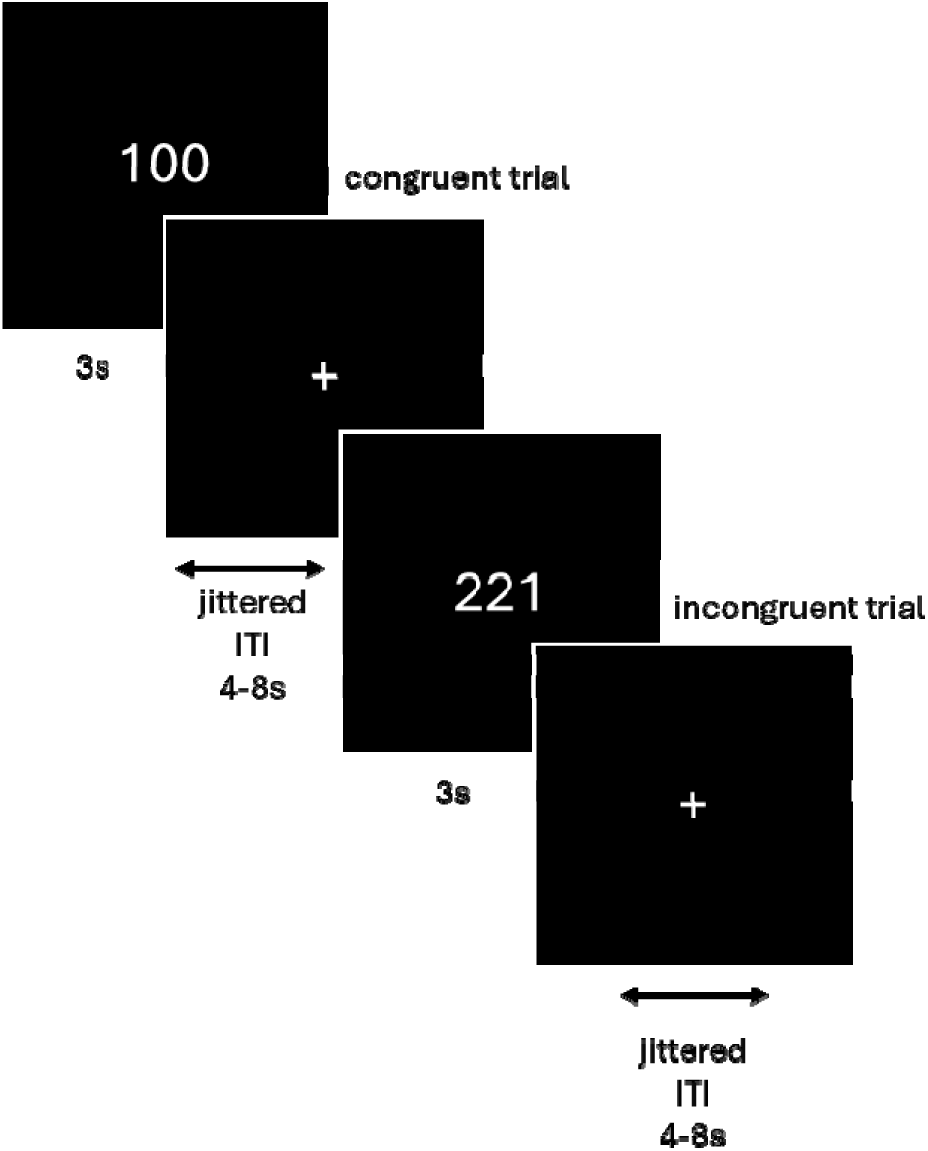
Illustration of MSIT trials. Congruent and incongruent trials were randomly presented using an event related design, with inter trial intervals jittered between target trials. Congruent trials presented the target number in a spatial position corresponding with the number. Incongruent trials presented target numbers in a spatial position in conflict with its position on the screen.

### MRI Acquisition

MRI scans were conducted using a Siemens Trio 3T scanner with a 12 channel RF coil. For high-resolution anatomical scan, a 3D magnetization-prepared rapid gradient-echo sequence (MPRAGE) T1-weighted sequence was used with the following parameters: TR=1.9s, TE=3.51ms, slices=208, matrix=192×256, flip angle 9°, resolution 1.0×1.0×1.0 mm^3^. For functional imaging including a 8-minute resting state and a 11-minute task, blood-oxygen-level-dependent (BOLD) fMRI scan, a gradient-echo, echo-planar images (EPI) sequence was used with an oblique axial scans 30° from the AC-PC line and AP phase encoding direction and the following parameters: TR=2s, TE=27ms, flip=78°, voxel resolution=3.4375×3.4375×4 mm.

### Data Analyses

#### Resting state fMRI coactivation pattern analysis (CAPS)

Here we applied Co-Activation Pattern Analysis (CAPs) to assess dynamic resting-state brain activity, following the approach validated by Janes, et al. ^25^, in which eight transient brain dynamic states were derived that align with previously established resting state networks: frontoinsular-DMN, FPN, DMN, DAN, salience network (SN), occipital sensory-motor (OSM), DMN-OSM, and SN-O. In the current study we indexed network temporal dynamics using the Capcalc package (https://github.com/bbfre derick/capcalc). The parameter for total time in each state was derived (time spent in each state across the entire resting-state scan). Full results of dynamic resting-state brain activity for all 8 states under placebo for this sample are reported in our prior study [27]. In the current study, we focused on the time spent in the DAN state during the placebo session, due to its sensitivity to dopaminergic modulation and to reduce the space of tested variables. See Figure 2 for a visualization of the DAN network state.

**Figure 2.**
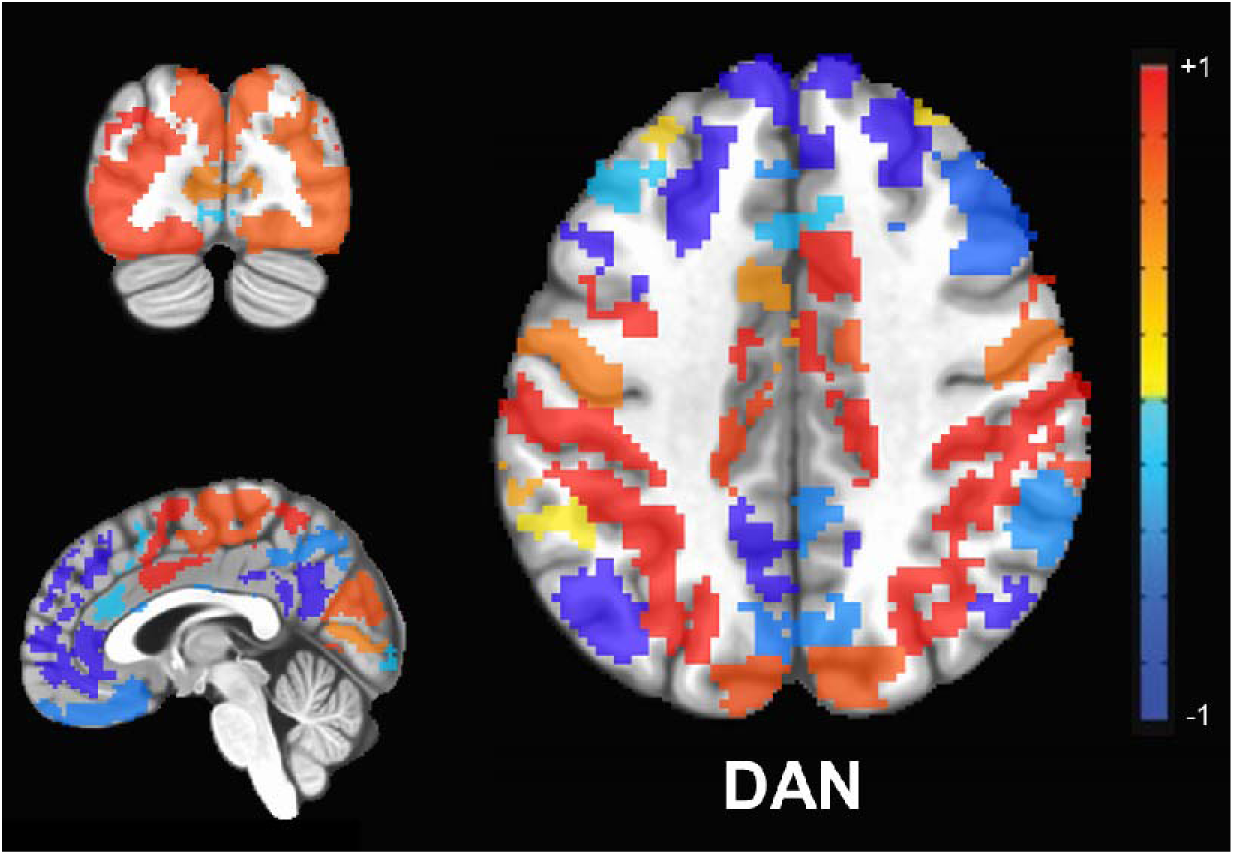
Visualization of DAN state derived using CAPs. Time spent in the DAN state under placebo was used to determine the relationship between dynamic brain activity at rest and task-based activation on the MSIT after participants had received methylphenidate or haloperidol. Warm colors denote areas of relative activation, while cool colors denote areas of relative deactivation. Reproduced with permission from Hill, et al. ^27^. DAN: Dorsal attention network.

**Figure 3.**
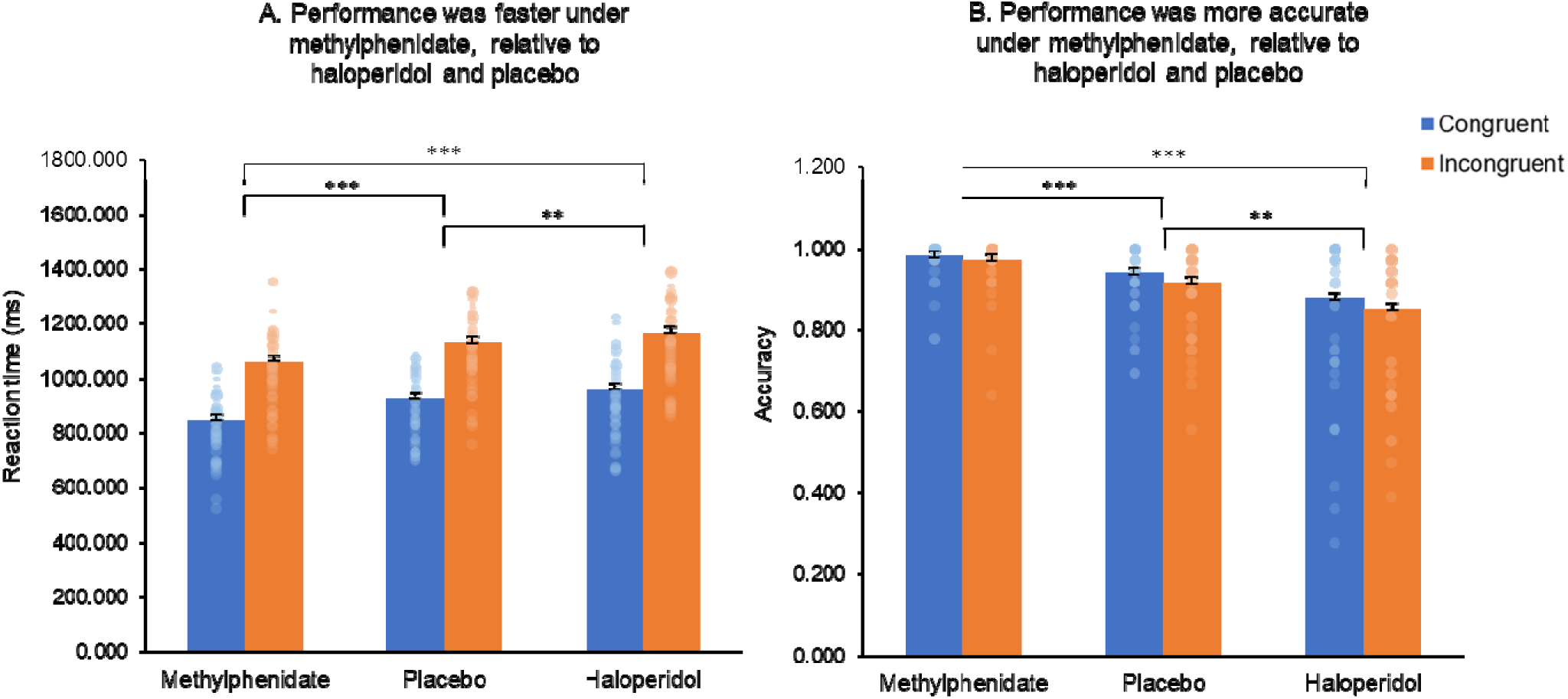
MSIT behavioral performance for (A) Average reaction time; and (B) Average accuracy. *** p<0.001; ** p<0.05

#### MSIT behavioral analyses

Analyses of behavioral metrics were conducted in SPSS Version 29. The behavioral data of four subjects were excluded due to poor performance on congruent trials in the placebo condition (values less than 3^rd^ quartile + 3 x interquartile range) and therefore were excluded from all subsequent analyses. On the remaining 55 participants, two separate within-subjects ANOVAs for 1) reaction time and 2) accuracy evaluated main effects of *drug condition* (methylphenidate, haloperidol, placebo) and *trial type* (congruent, incongruent), and their interaction.

#### MSIT task fMRI analyses

##### Preprocessing

The MRI DICOM data were converted to the BIDS compatible structure/format with bidskit (v2022.5.24a, https://github.com/jmtyszka/bidskit). The MRI preprocessing was conducted using fMRIPrep (v22.1.0, https://fmriprep.org/en/22.1.0/) and AFNI (v20.2.06, http://afni.nimh.nih.gov/afni/), which included discarding the first three volumes, slice timing correction, volume registration and six directional head motion parameter estimation, and spatial normalization to MNI space and resample to 2mm isotropic resolution. Head motion was also evaluated at the volume-by-volume level to further control image quality by using frame-wise displacement calculated with the Euclidean distance (*1d_tool.py*, AFNI). Volumes with displacement > 0.35 mm were censored (along with the volumes from the previous TR). The aCompCor was estimated to model physiological confound [53] by calculating the principal components of the signal from the white matter and cerebrospinal fluid (*3dmaskSVD*, AFNI). A set of polynomial terms was determined automatically based on the length of the fMRI time series for detrending (*3dDeconvolve*, AFNI), and a set of high-pass filter regressors was also generated via discrete sine/cosine transform with a cut-off frequency 0.01Hz (*1dBport*, AFNI). Four task-related GLM terms (congruent correct, incongruent correct, congruent incorrect, incongruent incorrect) were modeled along with all noise regressors with a single step of general linear model (GLM), pre-whitening to model the temporal autocorrelation during nuisance regression (*3dREMLfit*, AFNI) [54,55]. Finally, the data was smoothed with 6mm FWHM for subsequent group level analyses (*3dmerge*, AFNI).

##### Relationship between task activation and resting dynamic index

Group level analyses were conducted with linear mixed effect modeling (*3dLME*, AFNI) to probe the relationship between MSIT task activation under different drug conditions (methylphenidate, haloperidol) and the time spent in DAN state under placebo. The main effect time in DAN under placebo, main effect of drug, and their interaction (time in DAN x drug) were included in the model: MSIT activation ∼ drug x time in DAN under placebo + age + drug x age + sex + drug x sex + mean head motion + drug x head motion (1|Subject). Task activation was defined as the incongruent vs. congruent contrast for correct trials. Age, sex and mean head motion, as well as their interactions with the drug condition, were included in the model as covariates. Statistical significance was determined at voxel-level p<0.001 and Monte-Carlo simulation based multiple comparison correction at cluster level α<0.05, NN1 (face-wise nearest neighbor), (*3dClustSim*, AFNI). The beta values of the significant ROI identified by the interaction were examined and one outlier was identified as having a value greater than 3rd quartile + 3 x interquartile range. This subject was subsequently excluded from group level analyses, leaving 54 subjects for modelling the relationship between the dynamic resting index and task activation.

## Results

### Behavioral MSIT results

#### Reaction time

A *drug condition x trial type* ANOVA revealed a significant main effect of drug (*F*_2,108_ = 21.407, *p* < 0.001) and trial type (*F*_1,54_ = 418.636, *p* < 0.001) but no interaction (*p* = 0.824). Subjects performed faster for congruent (910.759ms ± 153.490ms) relative to incongruent trials (1119.031ms ± 170.577ms). Participants responded faster under methylphenidate (953.667ms ± 184.794ms) relative to haloperidol (1063.009ms ± 193.960ms; *p* < 0.001) or placebo (1028.008ms ± 184.374ms; *p* < 0.001). Participants also responded faster under placebo relative to haloperidol (*p* = 0.018). See Figure 2A.

#### Accuracy

A *drug condition x trial type* ANOVA revealed a main effect of drug (*F*_2,108_ = 22.637, *p* < 0.001) and trial type (*F*_1,54_ = 17.445, *p* < 0.001) but no interaction (*p* = 0.253). Participants were more accurate for congruent (0.935 ± 0.125) relative to incongruent trials (0.913 ± 0.125). Participants performed more accurately under methylphenidate (0.977 ± 0.056) relative to haloperidol (0.865 ± 0.171) or placebo (0.929 ± 0.094). Participants were also more accurate under placebo relative to haloperidol (*p* < 0.001). See Figure 2B.

### Time in DAN and MSIT task activation results

A linear mixed effects model revealed a significant interaction between time spent in the DAN under placebo and task-based activation during the MSIT under haloperidol/methylphenidate in the dorsolateral prefrontal cortex (dlPFC), peak MNI coordinates: [-43, 45, 1]; cluster size = 424mm^3^ (53 voxels x 8 mm), p < 0.001; cluster size cutoff = 360mm^3^ (45 voxels x 8 mm) for MSIT activation (see Figure 4). Post-hoc analyses indicated that there was a positive correlation between time spent in DAN at rest and dlPFC activation for incongruent vs. congruent MSIT trials under haloperidol (*r* = 0.471) and negative correlation under methylphenidate (*r* = −0.424; see Figure 5). Analyses controlled for age, sex and head motion. There was no significant main effect of drug on MSIT activation. There were also no significant correlations between dlPFC activation and RT or accuracy on the MSIT under methylphenidate or haloperidol (all *p*’s > 0.05). The whole brain activation pattern for the incongruent – congruent contrast for haloperidol, methylphenidate and placebo displays the expected pattern of activation [32,56] across all drug conditions. Activation maps and beta values for the spread of dlPFC activation are presented in the Supplementary Material (see Figures S2 and S3).

**Figure 4.**
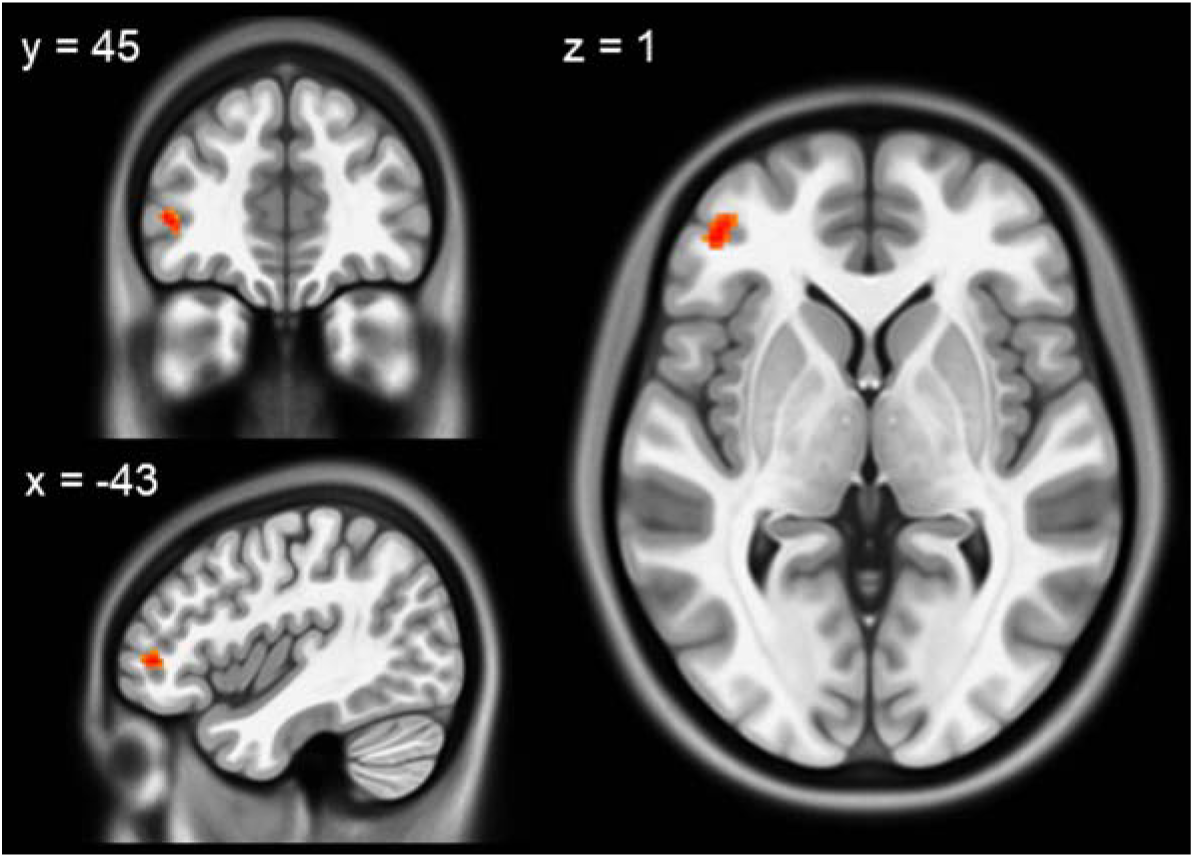
Interaction between DAN resting state dynamics and drug. A linear mixed effects model of time spent in the DAN at rest under placebo and task activation for the contrast of incongruent - congruent trials during the MSIT indicated a significant interaction in the dlPFC.

**Figure 5.**
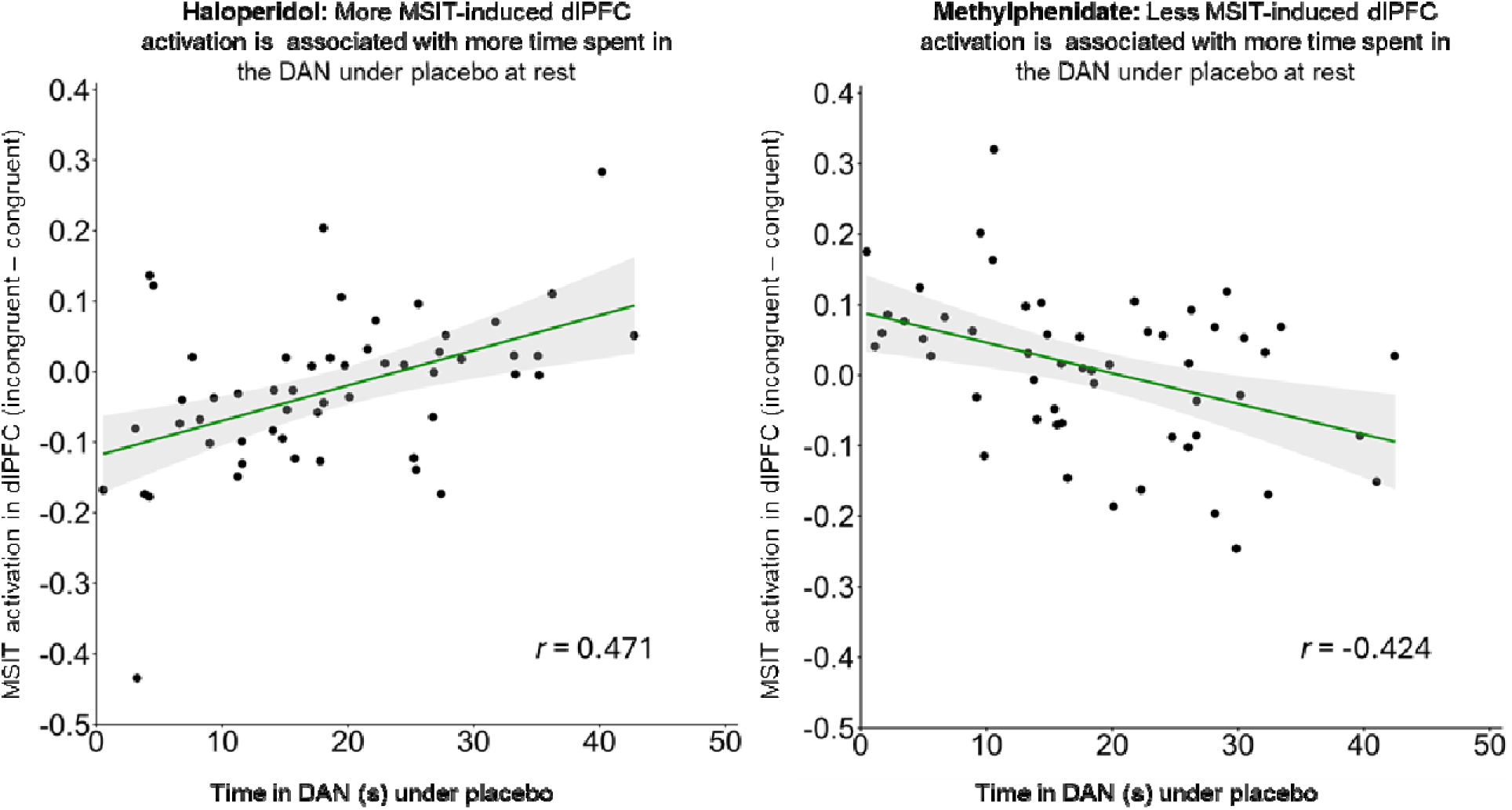
Post-hoc analyses of resting state dynamics and drug interaction. Time in DAN under placebo was associated with dlPFC activation during the MSIT in a drug-specific manner, where the relationship was positive under A) haloperidol; and negative under B) methylphenidate. DAN: dorsal attention network.

## Discussion

The current findings show that individual differences in resting-state brain dynamics under placebo are linked to variability in task-related brain activation across different drug conditions. Specifically, we found that the interaction between resting-state dynamics of the dorsal attention network (DAN) under placebo and the pharmacological condition was associated with task-related activation in the dorsolateral prefrontal cortex (dlPFC). Post hoc analyses revealed a drug-specific relationship: greater time spent in the DAN at rest under placebo was associated with increased dlPFC activation under haloperidol but decreased activation under methylphenidate in response to demands for external attention on the MSIT. This suggests that drug-free resting brain function may be a useful signature of cognitive processes influenced by pharmacology.

Here we found that the dlPFC was the site of interaction between drug state and time spent in the DAN at rest. The identification of the dlPFC in the context of the MSIT, fits with previous literature, as this region is consistently implicated in tasks requiring top-down attentional control [57–60], including the MSIT [32,61]. The dlPFC is also a central hub within the frontoparietal network [FPN; 62]. The FPN supports goal directed cognition and facilitates performance on externally directed cognitive tasks by coupling with the DAN [63], including on the MSIT [33]. While the dlPFC is typically activated alongside other regions within the FPN during cognitive tasks [62], in our study, task-based activation was limited to the dlPFC, rather than the entire FPN, and this activation was modulated by drug state. These current findings fit with the literature as engagement of the dlPFC is sensitive to variance in dopamine and norepinephrine function [64,65]. Previous work shows that haloperidol dampens prefrontal dopamine signaling [66], which likely contributes to its impairing effects on cognitive processing [67]. In contrast, methylphenidate enhances catecholaminergic neurotransmission in the prefrontal cortex [68], which may underlie its role in improving cognitive performance [69]. At the group level, our results are consistent with these behavioral findings, where performance on the MSIT was faster and more accurate under methylphenidate and slower and less accurate under haloperidol (however these responses were not orthogonal within individuals, see supplementary results). These opposing effects likely reflect drug-induced modulation of dlPFC function and help to explain the distinct patterns of dlPFC activation observed. The contrasting effects on prefrontal catecholamines thus offer a plausible neurochemical mechanism for the interaction between drug condition and time spent in the DAN within the dlPFC. Our results therefore suggest that individual variability in intrinsic brain dynamics helps shape variability during drug-related cognitive processes.

The current findings provide new evidence that individual differences in resting-state DAN dynamics predict how the dlPFC responds to pharmacological modulation during cognitive tasks, establishing a mechanistic link between intrinsic brain activity, catecholaminergic signaling, and inter-individual variability in drug response. Although the precise neurobiological mechanisms underlying DAN temporal dynamics remain unclear, emerging evidence suggests a role for catecholaminergic signaling. The DAN overlaps with regions rich in dopamine transporters [52], and both dopamine and norepinephrine function have been linked to DAN connectivity [43] and activity [27,40]. Our recent work shows that methylphenidate increases time spent in the DAN at rest [27], providing direct evidence that DAN temporal dynamics are sensitive to catecholaminergic modulation. These findings indicate that individual differences in resting-state DAN dynamics may partially reflect variation in underlying catecholaminergic function, which could in turn influence how regions such as the dlPFC respond during cognitive tasks under different pharmacological conditions. This interpretation is supported by Bush et al. [70], who demonstrated that methylphenidate increases dlPFC activation during the MSIT, highlighting this region’s sensitivity to catecholaminergic modulation. While further research is needed to definitively establish a causal link between DAN dynamics and catecholaminergic signaling, the current study advances the field by moving beyond descriptive observations of variability. Specifically, it demonstrates that resting-state temporal dynamics interact with pharmacological state to influence task-evoked activation, providing a framework to understand inter-individual differences in cognitive and pharmacological responsiveness.

Rather than correlating resting-state properties with subsequent task function [1,6], the current work took an exploratory approach to identify brain regions where task-evoked activation reflected an interaction between drug-free temporal dynamics at rest and pharmacological state. As such, the brain region identified by this interaction analysis (i.e., the dlPFC) fell outside the regions typically found to be significant during the standard MSIT incongruent-congruent contrast. Other work has also found that methylphenidate and atomoxetine increase dlPFC during the MSIT [70,71]. Thus, the current study not only supports but also extends prior findings by demonstrating that variability in DAN dynamics is linked to changes in dlPFC activity when catecholaminergic function is altered pharmacologically. This exploratory approach therefore highlighted inter-subject variability in drug-specific dlPFC activation that was related to variability in DAN temporal dynamics at rest under placebo – a finding which may have otherwise been missed using more a-priori approaches. While future work may may use hypothesis-driven strategies with a focus on regions that are canonically activated during the MSIT, our work nevertheless provides a novel approach to understanding how temporal dynamics at rest contribute to task-based brain activation.

It should also be noted that other networks involved in cognition may also relate to task-based activation under the drug conditions. Extensive evidence indicates that both the FPN and the default mode network [DMN; 72] play key roles in cognitive function, and that interactions between the DAN and these networks support cognitive processes [33,73,74]. Our supplementary analyses show that time spent in the DAN at rest under placebo is positively correlated with time spent in the DMN and negatively with time spent in the FPN. Further analyses of the interactions between drug and time spent in the FPN or DMN on dlPFC activity, as defined by the DAN-focused analysis, mirrored the correlation patterns observed between the time spent in the DAN, DMN and FPN at rest (see Figure S4). This is perhaps unsurprising, given that the dlPFC region was defined by the DAN-focused analyses, and given the well-characterized relationships between the DAN, FPN and DMN. We may therefore anticipate that activity in these networks would also relate to dlPFC activation. Nevertheless, these supplemental analyses provide preliminary evidence that resting-state temporal dynamics of other cognitive networks may also relate to task-activation. However, to fully explore how other network activity interact with drug to influence task-based activation, studies with larger samples are needed.

Despite the strengths of our current findings, there are several limitations. As discussed above, the current work cannot directly establish a link between individual variability in catecholaminergic function and DAN dynamics. To establish this link, future work should employ PET imaging alongside fMRI. Moreover, future work assessing the temporal properties of resting state activity in clinical populations with known variance in dopaminergic profile, such as in those with ADHD, may also help to elucidate the mechanisms that link baseline DAN activity with task-based medication response, as well as assess the clinical applicability of this approach. There were also no correlations between behavior and activity in the dlPFC. Notably, previous work indicates that brain-behavior correlations are not necessary to observe task-based brain activation that supports MSIT performance [34,70]. Moreover, the fact that dlPFC engagement differed under differing drug conditions suggests that drug state played an important role in how the dlPFC was recruited to do the task accurately. Nevertheless, future work in larger samples or in samples with greater variability in both behavior and brain activation, or even a more cognitively challenging task may produce a stronger association between behavior and brain activation. Further, the current work focused on attentional processes with the MSIT and medications that targeted catecholaminergic neurotransmission. Assessing other cognitive domains or using medications that target different neurotransmitters may offer new insights into the broader mechanisms linking resting state dynamics to task-based brain function.

Our results show that individual differences DAN function at rest in a drug-free condition provide insight into variability in medication response. The present findings make a significant step forward toward showing that inherent brain function has the potential to be used clinically to predict treatment response, which may contribute to the development of personalized, neuroscience-based treatment strategies.

## Acknowledgements

This research was supported [in part] by the Intramural Research Program of the National Institutes of Health (NIH). The contributions of the NIH author(s) are considered Works of the United States Government. The findings and conclusions presented in this paper are those of the author(s) and do not necessarily reflect the views of the NIH or the U.S. Department of Health and Human Services.

## Author contributions

KB drafted the initial manuscript. TZ and JH conducted data analysis. AJ, TZ and KB interpreted data. BJS contributed to study conceptualization and design. All authors critically revised the manuscript and approved the final version for submission.

## Data Availability Statement

The data that support the findings of this study are available from the corresponding author upon reasonable request.

