## Supplementary material for "From rest to focus: Pharmacological modulation of the relationship between resting state dynamics and task-based brain activation"

Clinical Trial Name & Number: Brain Networks and Addiction Susceptibility, NCT01924468

Exclusion criteria included, but were not limited to, a history of psychiatric disorders, HIV-positive status, or neurological illnesses. Medications that led to exclusion included, but were not limited to, benzodiazepines, barbiturates, anticonvulsants, antipsychotics, antidepressants, cold medications, and certain herbal supplements (e.g., Kava, Gingko biloba). All procedures were completed at the National Institute on Drug Abuse in Baltimore, Maryland and were approved by the institutional review board of the National Institutes of Health. Accrual began in August of 2013 and the study completed in September of 2018.

Sample size was determined by power analysis with the key power analysis pertaining to genotypic and fMRI data – target accrual assured that analysis fall into range to detect expected signal changes when accounting for intra-participant variance. This equaled 80 participants total – with 86 actual enrollees to account for drop out. Notably, this sample size was based in part by the genotypic analysis which was not of interest to the present study. The present sample size (N=59) is within standard range for the present study question using fMRI data.

This work was part of a larger trial where the primary goal was evaluating how genotype and catecholaminergic function relates to brain network activity during task and rest. This was assessed by measuring genotype information and measuring how modulation of the dopaminergic and nicotinic system via pharmacological manipulation alters brain network activity at rest and during cognitive tasks. The field has since largely moved past the genotypic analysis which informed the original primary objective, while new fMRI techniques have emerged for understanding brain activity, including coactivation pattern analysis to examine temporal dynamics of resting state brain activity. Here, we focused on the relationship between temporal dynamics of resting state activity, as measured by coactivation pattern analysis, and pharmacologically modulated task-based brain activity.

**Study day procedures**

All participants underwent scanning across three drug conditions, which were administered on separate days in a double-blind placebo-controlled design: placebo/placebo, placebo/methylphenidate, and placebo/haloperidol (see Supplementary Figure 1). The drug doses were 20 mg oral methylphenidate and 2mg oral haloperidol. Each scanning session was identical and took place at the peak drug effect: 3 hours post-haloperidol/placebo and 1 hour post-placebo/methylphenidate, timed according to absorption rates to ensure high and stable plasma levels of medication during the scan. To maintain blinding, participants received either drug or placebo twice on a given scan day—once at the haloperidol timepoint and once at the methylphenidate time point. On the placebo scan day, both timepoints involved the administration of placebo. Participants only received one active drug on each study day and placebo at the other timepoint. If participants were discontinued, slots were reused to maintain the balanced randomization. The study physician generated this randomization and assigned participants to interventions. The study physician and lead associate investigators were responsible for enrolling participants. All 59 subjects included in our analysis completed all sessions and did not having missing data.

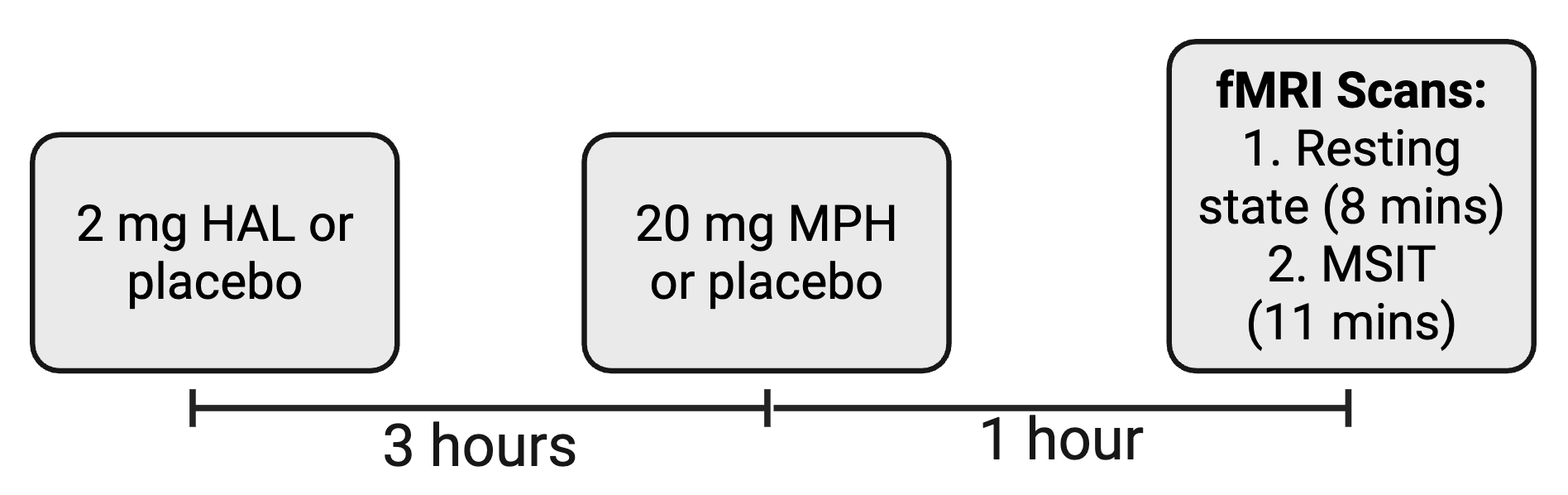

*Supplementary Figure 1. Study day procedures. Subjects received haloperidol, methylphenidate, or placebo timed in accordance with absorption rate to ensure drug peak coincided with the onset of the scanning session. HAL: haloperidol. MPH: methylphenidate.*

**Supplementary results**

*Behavioral results*

To measure whether there was an association between individuals who demonstrated an increase in reaction time (RT) and/or accuracy due to methylphenidate and those who demonstrated a reduction with haloperidol, we calculated the difference scores for RT and accuracy for methylphenidate-minus-placebo and haloperidol-minus-placebo and correlated these scores. There were no significant correlations between the difference scores for RT (*r* = 0.20, *p* = 0.15) or accuracy (*r* = 0.12, *p* = 0.38), indicating that there were no orthogonal responses to these medications within a subject.

*fMRI results*

Head motion (framewise displacement) differed significantly between methylphenidate (*M* = 0.069, *SD* = 0.024) and haloperidol (*M* = 0.091, *SD* = 0.028) conditions on the MSIT (*t*_53_ = 6.227, *p* < 0.001). The number of censored volumes also significantly differed between methylphenidate (*M* = 5.278, *SD* = 7.575) and haloperidol (*M* = 18.611, *SD* = 25.044) conditions on the MSIT (*t*_53_ = 3.782, *p* < 0.001). To control for this group difference, head motion was included in the model: MSIT activation ~ drug x time in DAN under placebo + age + drug x age + sex + drug x sex + mean head motion + drug x head motion (1|Subject).

Voxel-wise one sample t-tests (3dttest++, AFNI) were conducted for the incongruent-congruent contrast across drug conditions. Statistical significance was determined at voxel-level p<0.001 and Monte-Carlo simulation based multiple comparison correction at cluster level α<0.05, NN1 (face-wise nearest neighbor). The activation maps for the contrasts for methylphenidate, haloperidol and placebo are illustrated in Supplementary Figure 2.

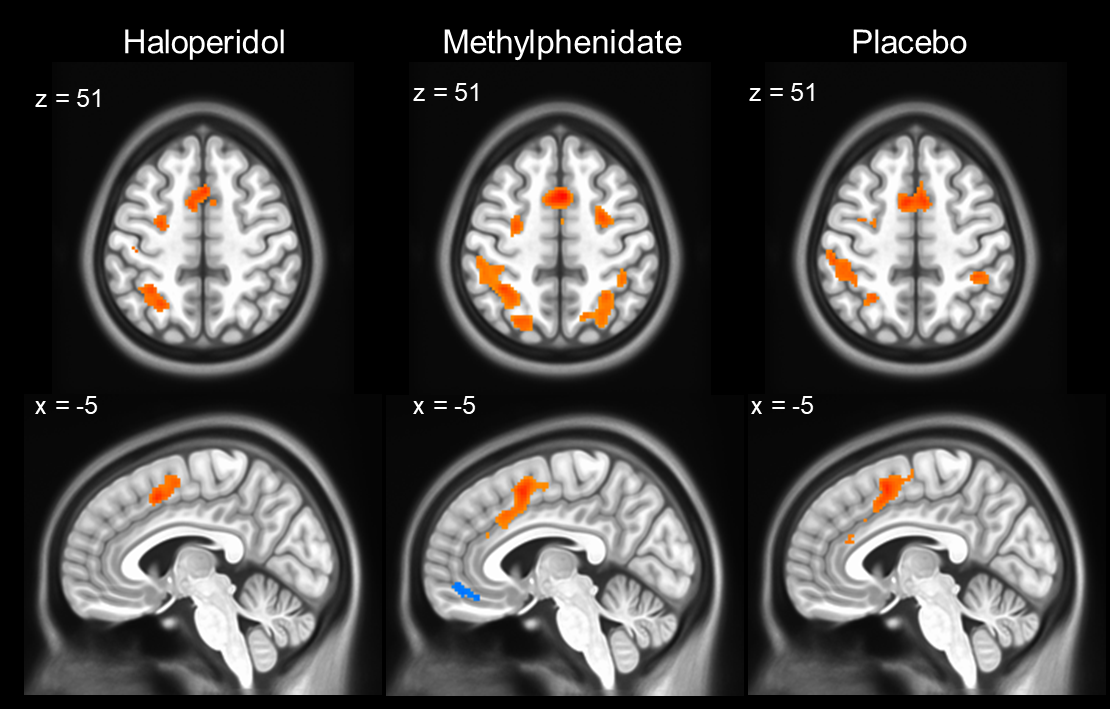

*Supplementary Figure 2. Activation map for MSIT task activation split by drug condition.*

Beta values for the spread of dlPFC activation for the interaction between time spent in the DAN under placebo and drug are presented in Supplementary Figure 3.

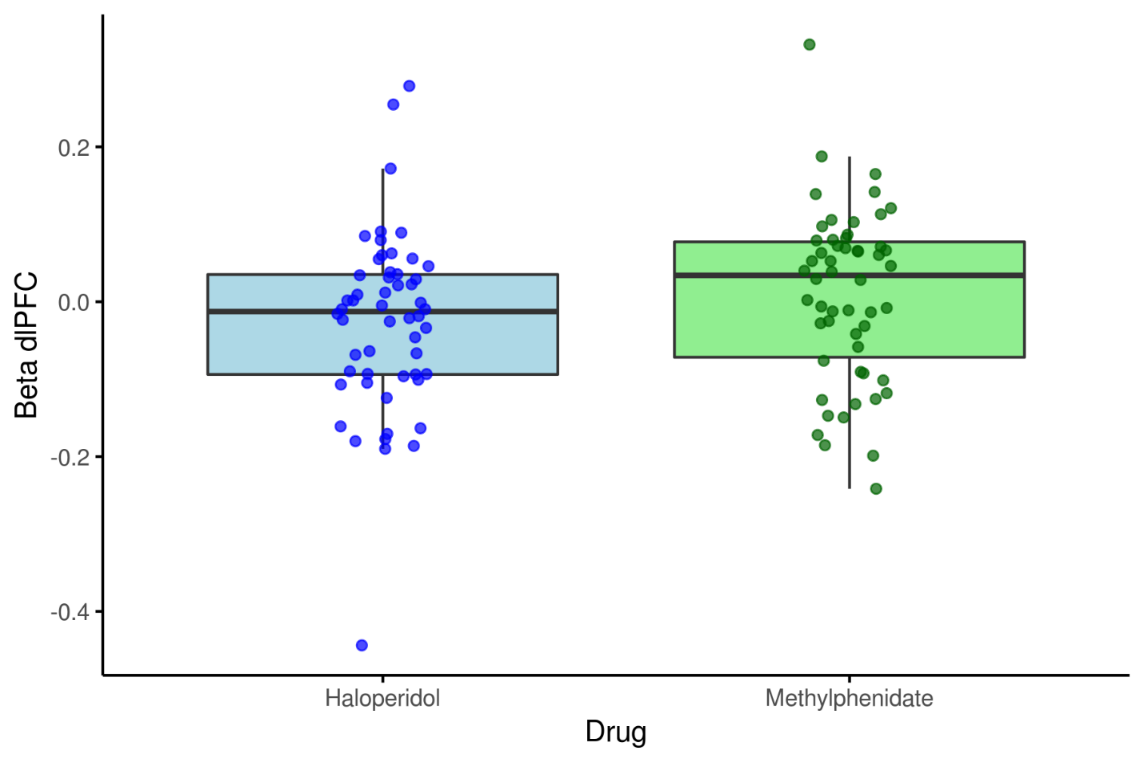

*Supplementary Figure 3. Spread of dlPFC activation (beta values) by drug.*

*Additional models of other time spent in default mode network (DMN) and frontoparietal network (FPN) under placebo at rest*

There were significant negative correlations between time spent in the DAN and FPN under placebo (*r* = -0.608, *p* < 0.001) and the FPN and DMN (*r* = -0.545, *p* < 0.01), while there was a significant positive correlation between time spent in the DAN and DMN under placebo (*r* = 0.672, *p* < 0.001).

Additional separate linear models of the interaction of time spent in the DMN and frontoparietal network FPN by drug in the dlPFC were both significant: drug x FPN (*F* = 19.30, *p* < 0.0001); drug x DMN (*F* = 5.82, *p* = 0.02). Post-hoc analyses of the interaction of each brain state by drug are illustrated in Supplementary Figure 4.

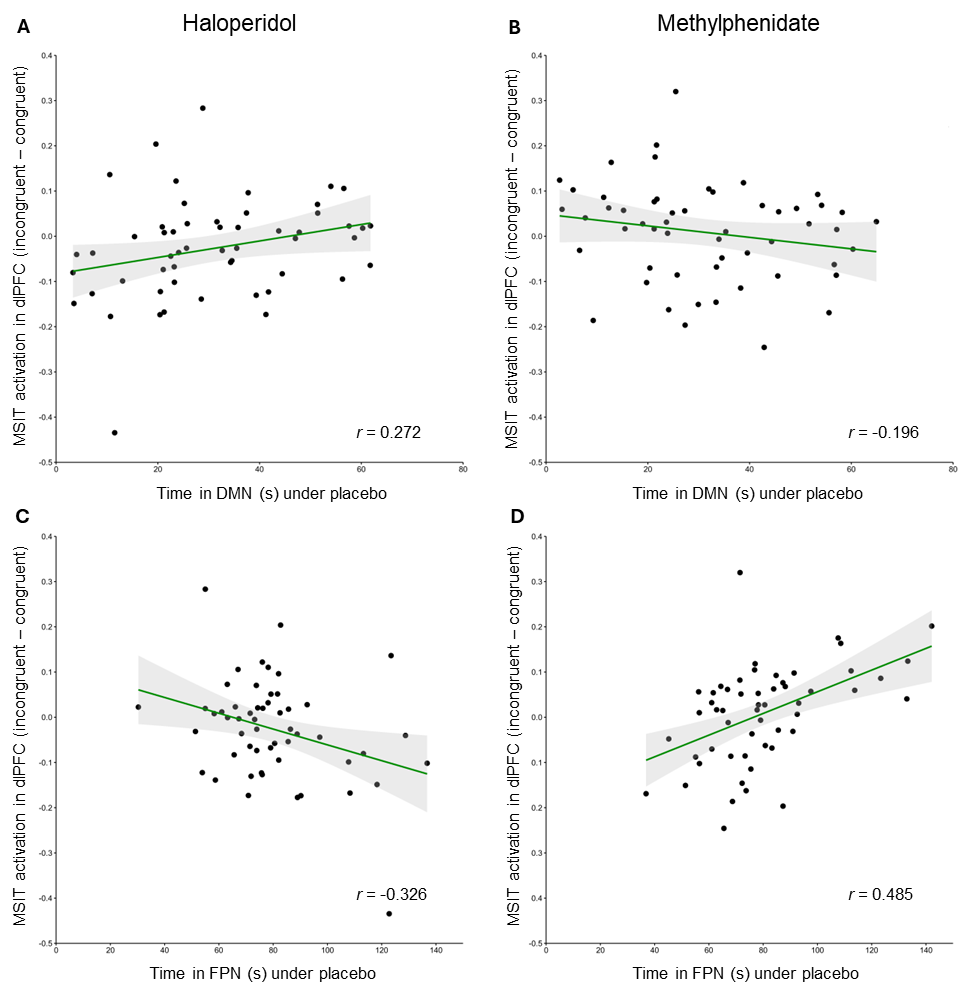

*Supplementary Figure 4. Post-hoc analyses of the interaction of time spent in the DMN and FPN under placebo and drug in the dlPFC during the MSIT. With the DMN the relationship was positive under A) haloperidol and negative under B) methylphenidate. With the FPN, the relationship was negative under C) haloperidol and positive under D) methylphenidate. dlPFC: dorsolateral prefrontal cortex; DMN: default mode network; FPN: frontoparietal network.*
